# Short chain fatty acids enhance expression and activity of the umami taste receptor in enteroendocrine cells via a Gα_i/o_ pathway

**DOI:** 10.1101/2020.06.01.127316

**Authors:** Matilda Shackley, Edward W. Tate, Alastair J.H. Brown, Gary Frost, Aylin C. Hanyaloglu

## Abstract

The short chain fatty acids (SCFAs) acetate, butyrate and propionate, are produced by the fermentation of non-digestible carbohydrates by the gut microbiota. SCFAs are of interest because they regulate appetite, adiposity, metabolism, glycemic control and immunity. SCFAs act at two distinct G protein-coupled receptors (GPCRs), FFAR2 and FFAR3. These are expressed in intestinal enteroendocrine cells (EECs), where they mediate SCFA-driven anorectic gut hormone release. EECs also express other GPCRs that act as nutrient sensors, in a manner that is plastic and adaptable to the environment. SCFAs may elicit some of their health-promoting effects by altering levels of GPCRs in EECs, thus, enhancing gut sensitivity to dietary molecules. Here, we identified that exposure of the murine EEC STC-1 cell-line to a concentration of SCFAs found in the colon, specifically enhances mRNA levels of the umami taste receptors TASR1 and TASR3, without altering levels of the SCFA GPCRs, FFAR2 and FFAR3. Interestingly, treatment of EECs with propionate or butyrate, but not acetate, increased levels of umami receptor transcripts. This phenomenon was reversed by inhibiting Gα_i/o_ signaling with pertussis toxin, suggesting that SCFAs act through FFAR2/3 to alter gene expression. Surprisingly, neither a FFAR3-nor a FFAR2-selective synthetic ligand could increase TASR1/TASR3 mRNA levels. We assessed the functional impact of increases in TASR1/TASR3 expression using unique pharmacological properties of the umami taste receptor; namely, the potentiation of signaling by inosine monophosphate. We found that the umami taste receptor induced inosine-1-phosphate and calcium signaling in response to L-alanine and L-monosodium glutamate, and that butyrate pretreatment significantly enhanced such signaling. Our study reveals that SCFAs may contribute to EEC adaptation and alter EEC sensitivity to bioactive nutrients.

## Introduction

After ingestion, physical and chemical processes digest food into a large and dynamic array of metabolites within the gastrointestinal (GI) tract. The detection of these, via ‘nutrient sensing’ mechanisms, results in the secretion of over 20 different peptides from enteroendocrine cells (EECs) (1). Of particular note are colonic short-chain fatty acids (SCFAs), the anaerobic fermentation of non-digestible carbohydrates, components of high-fiber diets. These are carboxylic acids with fewer than six carbons (Cs), which can reach high luminal concentrations of 10^−1^ M (2),(3). 95% of the SCFAs produced in the GI tract are acetate (2Cs), propionate (3Cs) and butyrate (4Cs) (3,4). These SCFAs, in particular propionate, are currently of interest, not only due to their ability to regulate anorectic gut hormone release, but also to promote weight loss, reduce abdominal adiposity and improve insulin sensitivity (5–8).

A large range of luminally expressed cell surface proteins is responsible for nutrient sensing. A significant proportion of these are members of the superfamily of G protein-coupled receptors (GPCRs) (8). SCFAs activate two distinct GPCRs that are known to be expressed in EECs, FFAR2 and FFAR3 (8–13). When expressed in heterologous cells, these two receptors display differential potency for each SCFA, which also differs between human and mouse receptor orthologs, yet propionate is the most potent of the SCFAs at both murine receptors (13–15). FFAR2 and FFAR3 both activate Gα_i/o_ signaling, and FFAR2 can also signal via Gα_q/11_ to release calcium (Ca^2+^) from intracellular stores; a pathway associated with its role in inducing gut hormone secretion from human and mouse EECs (5,7,8,16–18). However, beyond regulating levels of gut hormone expression (19) and secretion (5,7,17,18), our understanding of the additional roles of FFAR2/3 in EECs is limited.

A variety of GPCRs act as nutrient sensors in EECs, each responding to a distinct range of macromolecules and metabolites.(8) As the GPCR expression in EECs is not static (20), one possibility is that nutrients can alter GPCR expression levels, adapting the sensitivity of the gut to other dietary molecules. There is evidence to support this; obese individuals have significantly different expression profiles of nutrient sensing GPCRs in their GI tract compared with lean controls, with significant gene expression changes in genes encoding GPCRs, such as umami taste receptor subunit TAS1R3 and long chain fatty acid receptor FFAR4 (21,22). Further studies have demonstrated that there are significant differences in the mRNA expression of long and short chain fatty acid GPCRs and gustatory receptors in obese mice compared with lean controls (23), which are altered significantly following gastric bypass surgery (20). Overall, this suggests plasticity in the expression of nutrient sensing receptors, enabling dynamic adaptation to environmental factors. It is unknown whether the recently reported health benefits of increased colonic concentrations of SCFAs, such as propionate (5,7,17), are partly mediated by an underlying mechanism that alters the ability of the gut to sense other dietary components/metabolites.

In this study, we demonstrate that exposure of EECs to SCFAs can increase the gene expression of a specific gustatory GPCR, the umami taste receptor, without altering the levels of SCFA receptors. This altered gene expression was mediated by propionate and butyrate via a Gα_i/o_ signaling pathway, supporting a SCFA-GPCR signaling mechanism. However, synthetic FFAR2-or FFAR3-selective ligands could not mimic this. The increased expression of umami taste receptor subunits by SCFAs resulted in enhanced signaling from this receptor.

## Materials and Methods

### Cell culture

STC-1 cells originate from enteroendocrine tumors in the duodenum of double transgenic mice.(24) This cell line was used for all experiments. STC-1 cells were cultured (95% O_2_; 5% CO_2_; 37°C) in Dulbecco’s modified Eagle’s Medium (DMEM) containing 4.5 g/L D-glucose, 4 mM L-Glutamine (Sigma), supplemented with 10% FBS (Sigma), 100 U/mL penicillin and 100 mg/L streptomycin (ThermoFisher; DMEM+/+).

### Ligand treatment

STC-1 cells were grown to 70–80% confluency before treatment with SCFAs. All SCFAs were stored in solid salt form (Sigma). Solutions (100 mM) were made fresh for every experiment by dissolving in DMEM+/+ for incubations <5 hrs and in serum-free media for incubations <5 hrs. 2-(4-chlorophenyl)-3-methyl-N-(thiazole-2-yl)butanamide (4-CMTB; Tocris) was used as a FFAR2-specific agonist and AR420626 (Cayman) was used as an FFAR-specific agonist, both at a working concentration of 10 μM.

### Quantitative-PCR

After incubations with SCFAs, TRIzol® Reagent (Life Technologies) was used to extract RNA from STC-1 cells. After purification, 1 μg of each RNA sample was treated with an RNAse inhibitor (ThermoFisher) and a DNase I treatment kit (Life Technologies). SuperScript IV Reverse Transcriptase kit (Life Technologies) was used to synthesize complimentary DNA (cDNA). qPCR was performed using SYBR-Green PCR Mastermix kit (ThermoFisher). Each reaction was run in triplicates and cDNA was replaced with nuclease-free water as a negative control. Reactions were performed using the ABI StepONE sequence system. The 2^-ΔΔCT^ method(25) was used for analysis of raw C_t_ values. Briefly, gene expression was normalized to the housekeeping gene β-actin, and values from treated cells were compared to the expression of untreated controls. All primer sequences used were purchased predesigned from Sigma Aldrich UK (sequences found in **Supplementary Table 1**). Serial dilution curves were performed to ensure primer efficacy of 90–110%.

### Measurement of intracellular cAMP

All cAMP assays were performed in serum-free DMEM (Sigma) supplemented with 3-isobutyl-1-methylxanthine (IBMX; Sigma) to inhibit phosphodiesterase cAMP degradation. cAMP concentrations were measured from cell lysates after cells were incubated for 5 minutes with synthetic agonists (10 μM) for FFAR2 (4-CMTB) or FFAR3 (AR420626) using the HTRF cAMP Dynamic 2 immunoassay kit (CisBio). Fluorescence was measured with a PHERAstar FSX plate reader (BMG Labtech) equipped with HTRF 337 optic module, with excitation at 340 nm and measurements of emission at 620 nm and 665 nm. cAMP levels were interpolated from an cAMP standard curve and normalized to protein concentration. All experiments were conducted in triplicate and repeated at least 3 times.

### Measurement of intracellular inosine-1-phosphate (IP_1)_

IP_1_ signaling assays were performed after incubation with SCFAs to evaluate the response of STC-1 cells to L-monosodium glutamate (L-MSG; Sigma) and L-Alanine (L-Ala; Sigma), selected due to their potency at the rodent umami taste receptor (26),(27). All reactions were performed in the presence and absence of inosine monophosphate (IMP, 2 mM) in serum-free DMEM (Sigma) supplemented with 50 mM LiCl (Sigma). After cells were treated with ligands (30 min), IP_1_ concentrations were measured from cell lysates using the HTRF IP-One immunoassay kit (CisBio). Fluorescence was measured and IP_1_ levels were quantified using the same methodology as the cAMP assay.

### Ca^2+^ mobilization

Intracellular levels of Ca^2+^ were measured using the Fluo-4AM Direct Calcium Assay Kit (Invitrogen). STC-1 cells were incubated with a 1:1 ratio of opti-MEM media (Sigma, UK) to calcium dye Fluo-4-AM Direct for 30 min at 37°C and for a further 30 min at room temperature. Cells were imaged using a Leica Confocal Microscope (20X dry objective; 488 nm excitation). Movies were recorded at 1 fps for 60 sec before addition of IMP/control (2 mM). After ensuring no calcium mobilization in response to IMP, ligands (L-Ala or L-MSG) were added and movies were recorded until the readout returned to basal levels. All conditions for each experiment were conducted in duplicate and repeated at least three times. The fluorescence intensity of each cell was quantified using the ImageJ plugin Time Series Analyzer. The maximal intensity was obtained from subtracting the average background intensity (recorded before ligand addition) for each cell and averaged across 20 cells per condition.

### Statistical analysis

Data is represented as the mean ± the standard error (SE). GraphPad Prism was used to determine significance (p<0.05), using unpaired Student’s t tests, One-way ANOVA with a Dunnett post-hoc, or Two-way ANOVA followed by a Bonferroni post-hoc test.

## Results

### A physiologically relevant concentration of SCFAs significantly increases the expression of taste receptor transcripts

A key aim of our study was to determine whether SCFA treatment of EECs would alter GPCRs previously demonstrated to be differentially expressed between obese and lean mice and humans (20–23), with a specific focus on the taste receptor GPCRs. Initially we confirmed that STC-1 cells expressed FFAR2, FFAR3, TAS1R1, TASR2, TAS1R3, the taste receptor-associated G-protein, α-gustducin, and two bitter taste receptors, TAS2R(108) and TAS2R(138), that were selected based on their potential involvement in bitter compound-induced calcium signaling (28). We detected transcripts for all these genes in STC-1 cells, albeit in varying amounts (**Figure 1A**), confirming this cell-line represented an appropriate model in which to study potential interactions between SCFA signaling and the gustatory signaling system.

**Figure 1.**
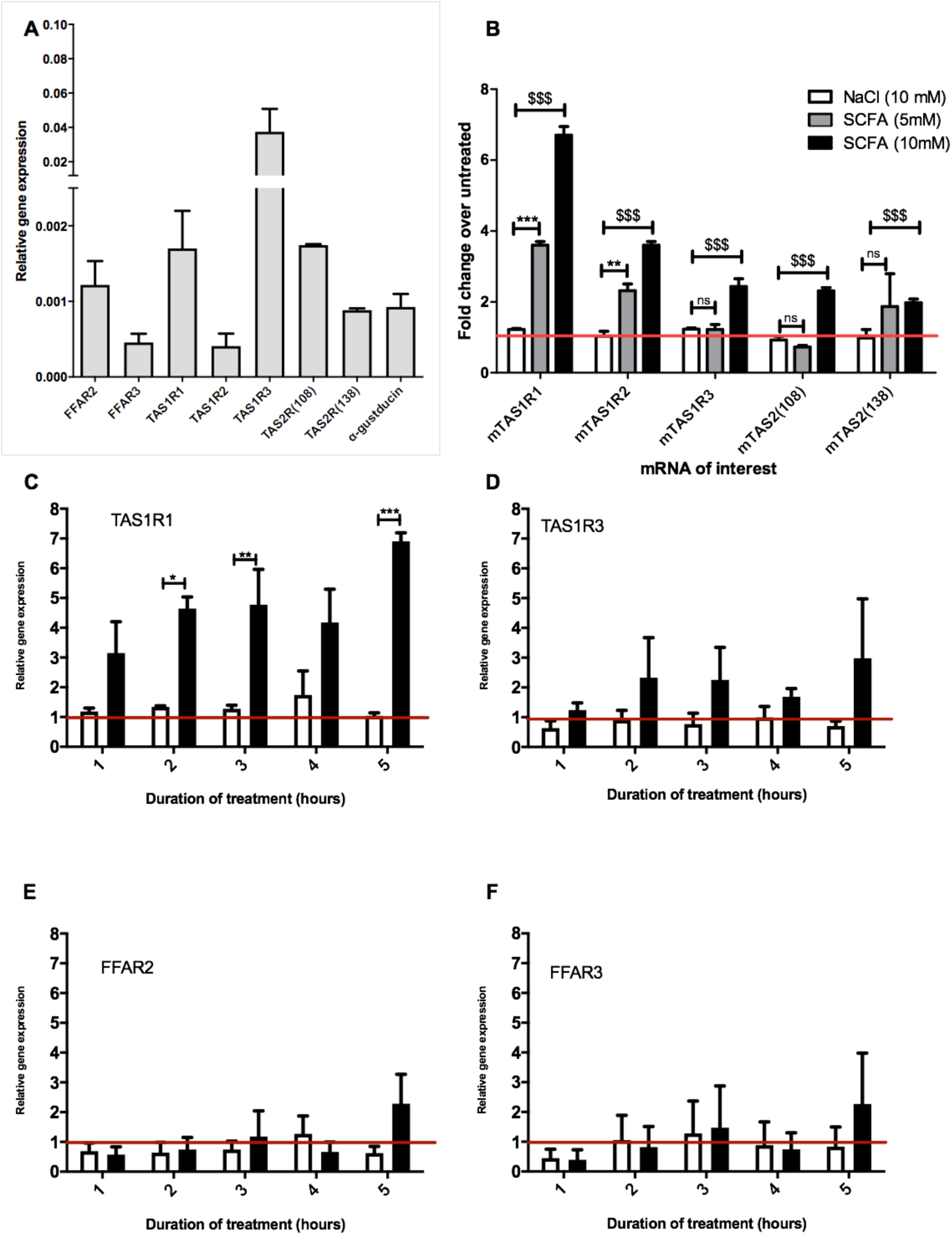
Exposure to SCFAs significantly changes the expression profile of taste receptors in STC-1 cells. **A**) RNA was extracted from STC-1 cells for qPCR analysis of taste receptors TAS1R1, TAS1R3, TAS1R2, TAS2(108) and TAS2(138); free fatty acids receptors FFAR2 and FFAR3; and taste-specific G-protein α-gustducin, and normalized to the levels of housekeeping gene β-actin. **B-G)** STC-1 cells were treated with NaCl (control; white bars), 5 mM SCFAs (grey bars) or 10 mM SCFA (black bars) for 2 hrs, after which RNA was extracted, purified and quantified with qPCR. Results are expressed as a fold change in expression over the untreated control and represents the average ± SEM, n=3. The line indicates a fold change of 1, where there has been no change in expression. Two-way ANOVA, with Bonferroni post hoc, $$$p<0.001 SCFA (10 mM) vs. NaCl control, **p<0.01, ***p<0.001 SCFA (5 mM) vs. NaCl control.

To determine whether SCFAs can influence the expression of taste GPCRs, STC-1 cells were incubated for 2 hrs with SCFAs in a 3:1:1 molar ratio of acetate:propionate:butyrate at 5 or 10mM (final concentration) chosen to reflect the physiological SCFA concentrations in the proximal and distal colon(3,4). qPCR was used to analyze the relative changes in expression of the transcripts of TAS1R1, TAS1R2, TAS1R2, TAS2 (108) and TAS2 (138). Incubation with 10 mM SCFAs significantly upregulated all taste receptors (p<0.001 vs. control), whereas incubation at 5 mM only significantly upregulated TAS1R1 and TAS1R2 (**Figure 1B**). The largest fold change was observed with transcripts for TAS1R1 where SCFAs (10 mM) induced a 6.7-fold increase over basal levels (**Figure 1B**). Based on these initial observations we decided to investigate the mechanism of upregulation of the TAS1R1 subunit further. As TAS1R1 is only functionally active when it is heterodimerized with TAS1R3 (the umami taste receptor),(26,27) we extended our investigation to include the TAS1R3 subunit. Treatment of cells with SCFAs (5 mM) over time (1–5 hr) revealed that TAS1R1 was significantly upregulated following 2, 3 and 5 hr of SCFA incubation (**Figure 1C**). Conversely, SCFAs at 5 mM did not affect the levels of TAS1R3 (**Figure 1D, 1B**). SCFAs did not alter the expression of SCFA receptors FFAR2 and FFAR3 (**Figure 1E, 1F**) at any time-point.

### The umami taste receptor is significantly upregulated by SCFAs, but not synthetic FFAR ligands

After demonstrating that a 3:1:1 mixture of SCFAs can influence the expression profiles of components of the umami taste receptor, we assessed whether specific SCFAs mediate these changes. STC-1 cells were treated with either acetate, propionate or butyrate (10 mM) for 5 hr, after which, mRNA levels of TAS1R1 and TAS1R3 were measured. Interestingly, incubation with propionate or butyrate, but not acetate, was sufficient to induce significant upregulation of both components of the umami taste receptor. TAS1R1 was upregulated ∼15-fold by both propionate (p=0.0186) and butyrate (p=0.0001; **Figure 2A**). TAS1R3 was upregulated more modestly than TAS1R1, by ∼3-fold following propionate (p=0.01) or butyrate (p=0.04) treatment (**Figure 2B**).

**Figure 2.**
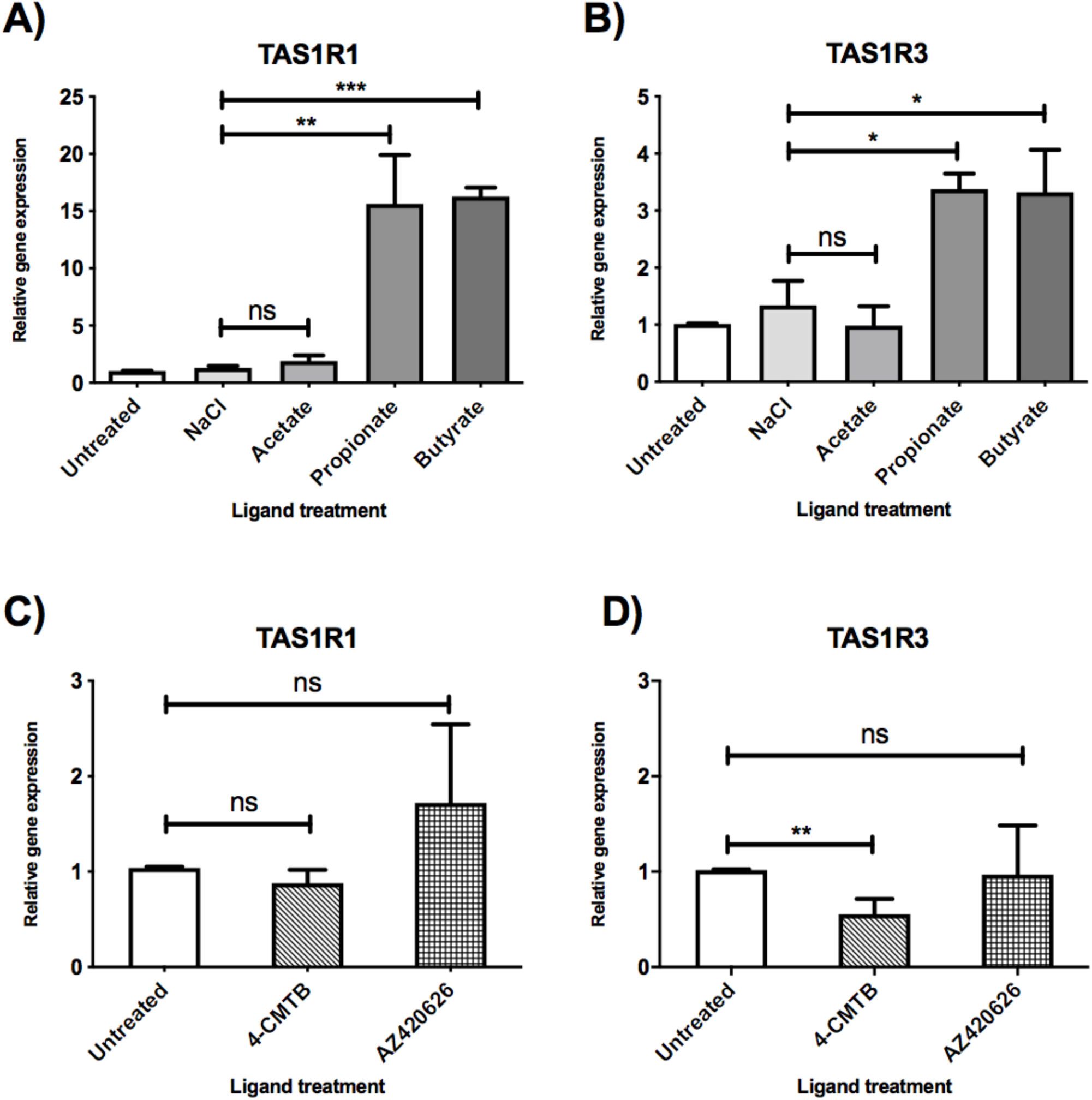
SCFAs and synthetic ligands differ in their ability to upregulate the umami taste receptor. SCFAs and FFAR2/3 agonists influence gene expression of TAS1R1 and TAS1R3 differentially in STC-1 cells **A-B**) STC-1 cells were incubated with NaCl or SCFAs (10 mM) for 5 hrs, after which, RNA was extracted and purified. Expression of taste receptors TAS1R1 (A), TAS1R3 (B) was quantified using qPCR analysis and normalized to the levels of housekeeping gene β-actin. Data are expressed as mean ± SEM fold-change in expression over the untreated control (n=3). T-tests vs. control; ns, non-significant; *p<0.05; **p<0.01; ***p<0.001. **C-D**) STC-1 cells were incubated with either 4-CMTB or AZ420626 (10 μM) for 5 hrs, after which, RNA was extracted and purified. Expression of taste receptors TAS1R1 (C), TAS1R3 (D) was quantified using qPCR analysis and normalized to the levels of housekeeping gene β-actin. Data are expressed as mean ± SEM fold change in expression over the untreated control (n=3). t-tests vs. control; ns, non-significant; *p<0.05; **p<0.01; ***p<0.001.

As SCFAs have been reported to be able to activate both FFAR2 and FFAR3 (9), we used synthetic ligands to determine whether selective activation of each receptor had a similar effect on umami taste receptor gene expression. STC-1 cells were exposed to 4-CMTB, a FFAR2-specific agonist, or AR420626, a FFAR3-specific agonist at concentrations known to induce maximal signal responses (12, 15). The ability of these synthetic ligands to activate the G*α*_i/o_ signaling, via inhibition of forskolin-induced increases in cAMP levels was also confirmed (See **Supplementary Figure 1**). While these ligands activate G*α*_i/o_ signaling, as do SCFAs, they were not able to upregulate the umami taste receptors (**Figure 2C, 2D**). Indeed, 4-CMTB induced a significant 2-fold decrease in mRNA levels of TAS1R3 (**Figure 2D**).

### SCFA-induced upregulation of the umami taste receptor involves Gα_i/o_ signaling

At rodent orthologs of FFAR2 and FFAR3, both propionate and butyrate show significant selectivity for FFAR3(15) a receptor known to signal via G*α*_i/o_ (9). To investigate whether G*α*_i/o_ activation plays a fundamental role in the SCFA-induced upregulation of umami taste receptor transcripts, STC-1 cells were incubated for 18 hrs with pertussis toxin (PTX), a Gα_i/o_ inhibitor. Compared to the basal levels of the PTX-pretreated control, pretreatment of cells with PTX significantly reduced the ability of propionate and butyrate to induce upregulation of TAS1R1, from 18.4-fold to 4.6-fold for propionate, and from 16.5-fold to 6.3-fold for butyrate (**Figure 3A**). PTX-pretreatment completely abolished the propionate- and butyrate-induced upregulation of TAS1R3 (**Figure 3B**).

**Figure 3.**
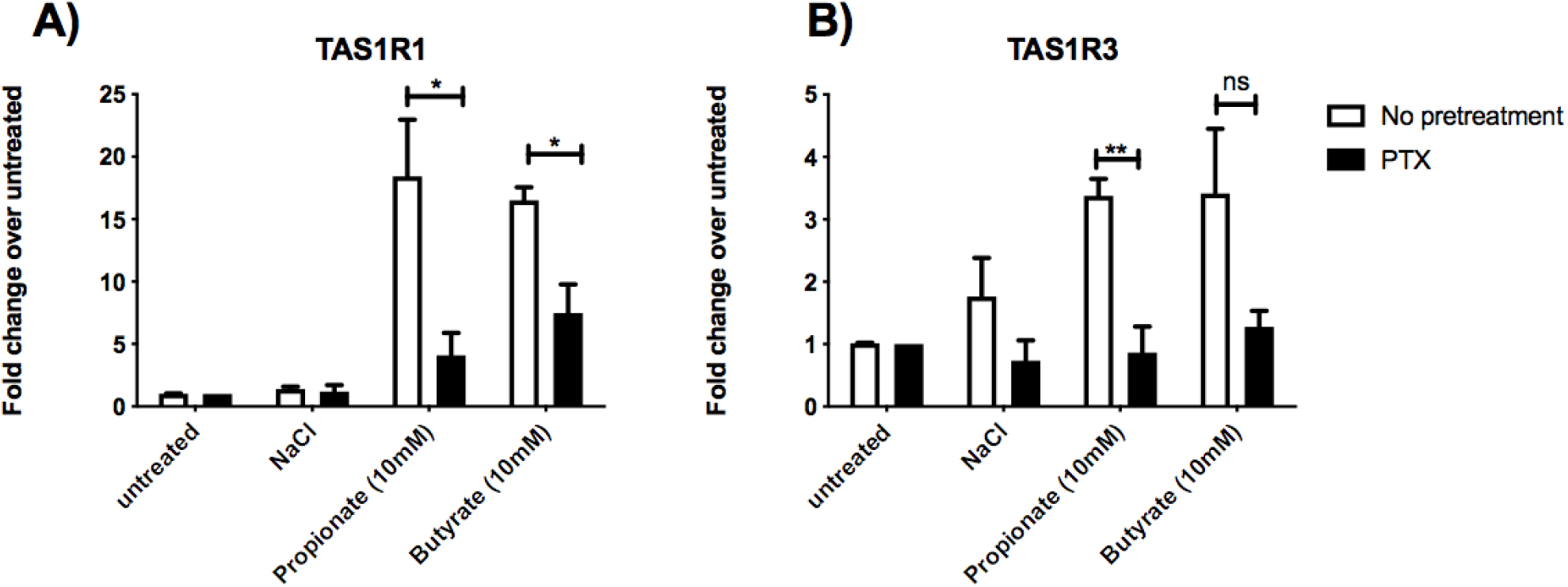
Inhibtion of Gα_i/o_ signaling impacts SCFA-mediated changes in taste receptor gene expression. **A-B)** STC-1 cells were pretreated with G*α*_i/o_ inhibitor pertussis toxin (PTX) (200 ng/μL, 18 hrs; black bars) or no pretreatment (white bars), followed by stimulation with either NaCl, propionate or butyrate (all 5 mM) for 5 hrs. RNA was extracted and purified. Expression of taste receptors TAS1R1 (A), TAS1R3 (B) was quantified using qPCR analysis and normalized to the levels of housekeeping gene β-actin. Data are expressed as mean ± SEM fold change in expression over the NaCl control either with or without PTX exposure (n=3). Two-way ANOVA, Bonferroni post hoc of no pretreatment vs. PTX treatment for each ligand; ns, non-significant; *p<0.05; **p<0.01; ***p<0.001.

### Umami taste ligands signal in STC-1 cells in a manner that is potentiated by addition of IMP

We then aimed to determine whether the observed upregulation of umami taste receptor mRNA could be translated into an increase in functional umami receptor signaling. The umami taste receptor is sensitive to a number of L-amino acids. L-MSG is the characteristic umami-tasting ligand, but studies have shown this to be less potent at the mouse isoform of the receptor than at the human.(27) It is documented that L-Ala elicits the strongest Ca^2+^ signals at the murine umami receptor.(27) Thus, we selected L-Ala for use in our assays. To confirm the signals were via activation of umami taste receptor, rather than other amino acid-sensitive receptors, we first assessed whether signaling was synergized by IMP, as this is a unique signaling property of the umami receptor.(26,27),(29) Taste receptors have been shown to activate phospholipase C-mediated pathways, leading to formation of 1,4,5-inositol triphosphate (IP_3_) (29), thus, umami taste receptor activation was determined by measurement of intracellular Ca^2+^and IP_1_, a downstream metabolite of IP_3._ Addition of IMP (2 mM) significantly increased the levels of Ca^2+^and IP_1_ signal induced by both L-MSG and L-Ala (10 mM; **Figure 4A, 4B**).

**Figure 4.**
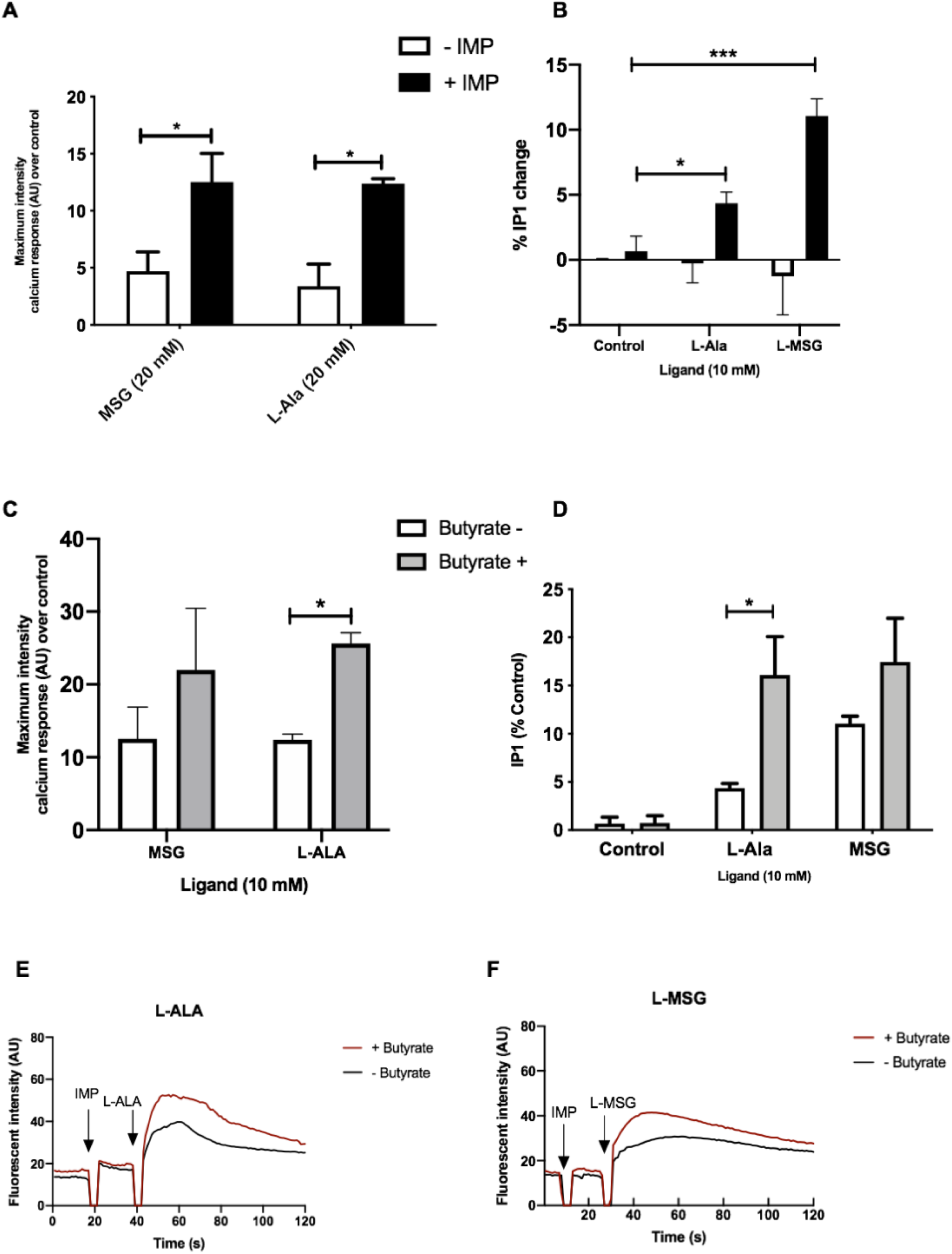
Umami receptor signaling cascades are potentiated by butyrate pretreatment. **A)** Intracellular calcium levels measured in STC-1 cells, incubated with fluorescence calcium indicator dye Fluo4-AM following addition of IMP (2 mM, black bars) or NaCl control (white bars; 2 mM) with L-Ala (20 mM) or L-MSG (20 mM). Data is expressed as mean ± SEM maximal fluorescence intensities over the control and is taken from 20 cells per sample, in duplicate (n=3). t-test vs. control; *p<0.05; **p<0.01; ***p<0.001. **B)** Intracellular IP_1_ accumulation measured in STC-1 cells on the addition of IMP (black bars) with either NaCl control (white bars; 2 mM), L-Ala (20 mM) or L-MSG (20 mM). Data is expressed as mean ± SEM across three distinct experiments; t-test *p<0.05; **p<0.01; ***p<0.001 vs. control. **C)** Intracellular calcium levels measured in butyrate-naive (white) or butyrate-pretreated (grey) STC-1 cells, incubated with fluorescence calcium indicator dye Fluo4-AM, followed by stimulation with IMP (2 mM) with L-Ala (20 mM) or L-MSG (20 mM). Data is expressed as mean ± SEM maximal fluorescence intensities over the control and are taken from 20 cells per sample, in duplicate (n=3). t-test vs. control, *p<0.05; **p<0.01; ***p<0.001. **D)** Intracellular IP_1_ accumulation measured in butyrate-naive (white) or butyrate-pretreated (grey) STC-1 cells after incubation with IMP (2 mM) and either L-Ala (20 mM) or L-MSG (20 mM). Data is expressed as mean ± SEM (n=3); t-test vs. control, *p<0.05; **p<0.01; ***p<0.001. **E-F)** Representative fluorescence intensity plots following IMP (2 mM) and L-Ala (E; 20 mM) or L-MSG (F; 20 mM) stimulation in butyrate pretreated (red lines) and butyrate naive (black lines) STC-1 cells, expressed in arbitrary units (AU).

### The increase in umami taste receptor transcript on exposure to SCFAs is coupled with an increase in signaling response to some umami taste ligands

To investigate whether upregulation of TAS1R1 and TAS1R3 mRNA by SCFAs translates into an increase in umami-receptor signaling, we pretreated STC-1 cells overnight with butyrate, at a concentration able to elicit significant upregulation of both transcripts (**Figure 2A, 2B**). We then reassessed the cells’ signaling response to L-MSG and L-Ala; pre-incubation with butyrate significantly increased IP_1_ signaling and the maximum-induced calcium response to L-Ala/IMP over time (**Figure 4C–F**). However, L-MSG-induced IP_1_ and Ca^2+^responses exhibited greater variability following butyrate pretreatment (**Figure 4C–F**), potentially consistent with the lower potency of this ligand compared to L-Ala at the rodent umami taste receptor (27). Overall, this data suggests butyrate-induced increases in umami taste receptor mRNA also result in enhanced umami taste receptor activity.

## Discussion

GPCRs expressed in the GI tract have a well-established role in nutrient-sensing and anorectic/incretin gut hormone secretion (3,5,7,8,16–19). Therefore, developing an understanding of GPCR expression profiles and signaling functions in EECs has therapeutic value in the field of obesity and Type II diabetes. SCFAs modulate gene expression in various cells, tissues and species (19,30–34). However, this is the first report that physiologically relevant concentrations of SCFAs, particularly propionate and butyrate, can directly and robustly upregulate transcripts encoding GPCRs in EECs. Of particular note, was the substantial upregulation of the umami taste receptor subunits, as the expression profile of these is significantly different in the GI tract of obese individuals when compared with lean controls.(20,21) These observations provide a mechanism to explain how diet composition and SCFA production are linked with fluctuations in GPCR expression patterns in obese humans and mice (20–23).

Our work demonstrated that the most highly upregulated taste receptor transcript upon EEC exposure to SCFAs was the umami taste receptor subunit TAS1R1. When exposed to a mixture of SCFAs, at a concentration often found in the colon (4,17), TAS1R1 was upregulated nearly 7-fold, without affecting expression levels of either of the SCFA receptors FFAR2/FFAR3. The umami taste receptor is a known heterodimer of TAS1R1 and TAS1R3 (26,27,29). It is co-expressed in GI tissue with CCK (35) and, on activation by protein hydrolysates, induces cholecystokinin (CCK) secretion from EECs (36). Interestingly, exposure to either propionate or butyrate robustly enhanced gene expression of both umami taste receptor subunits. There are two plausible mechanisms for the effects of SCFAs on the expression of umami receptor transcripts; through FFAR2/3 G-protein signaling or through histone deacetylase (HDAC) inhibition (37). That both propionate and butyrate, but not acetate, can increase the levels of these receptor is interesting, and maybe explained by the difference in potency and affinity of the SCFAs at rodent FFAR2/FFAR3 (15) and at HDACs (37).

Both FFAR2 and FFAR3 couple to G*α*_i/o_ signaling, and our data support a role for this GPCR signaling pathway in mediating the upregulation of taste receptor genes. We clarified the contribution of Gα_i/o_ signaling elicited by SCFAs (9,12,14,15) by inhibiting FFAR2/3 Gα_i/o_ signaling with PTX. PTX significantly reduced the upregulation of both umami receptor subunits induced by both propionate and butyrate. This suggests FFAR2/3 Gα_i/o_ signaling contributes significantly to the upregulation, even for butyrate; a very potent HDAC inhibitor (<1 mM) (37). Inhibition of G*α*_i/o_ activity abolished SCFA-induced TAS1R3 upregulation, and significantly reduced propionate-induced TAS1R1 upregulation more than butyrate-induced upregulation. Interestingly, propionate cannot inhibit HDACs as potently as butyrate, only doing so at high concentrations of >10 mM (37). Together, this potentially suggests that propionate acts more potently than butyrate through a FFAR2/3 signaling mechanism to induce TAS1R1 expression. If propionate is acting via FFAR2/3 to modulate gene expression, it may be hypothesized that FFAR3 is the more likely candidate, as propionate is nearly ten times more selective for rodent FFAR3 than FFAR2 (15). Furthermore, FFAR3 signaling influences gene expression in other cellular models: FFAR3 KO murine pancreatic islets have significantly different transcriptomes to wild-type animals, though in genes associated with insulin secretion and glucose regulation (38).

Surprisingly, synthetic FFAR2/FFAR3 selective ligands could not upregulate TAS1R1 or TAS1R3 transcripts, despite their ability to activate upstream receptor signaling in EECs. This suggests that endogenous SCFA and synthetic ligands have distinct activation profiles at FFAR2/3, thus, potentially eliciting different downstream responses. If both SCFAs and synthetic ligands activate similar upstream G-protein pathways, it remains to be determined the additional mechanisms that drive the SCFA-selective increases in gene transcription of umami taste receptors, but potentially suggests a role for ligand-induced bias signaling at FFAR2/3.

We then determined whether SCFAs could enhance functional umami taste receptor activity in STC-1 cells. Other GPCRs able to sense L-AAs are also expressed in EECs (8,39), but there were some technical challenges in deciphering the precise contributions of each L-AA-sensitive GPCR, owing to the lack of selective ligands. However, the synergistic effects of IMP offered a mechanism to detect umami-specific responses (26,27). Here, umami ligands, L-MSG and L-Ala, only induced increases in Ca^2+^and IP_1_ in the presence of IMP, as observed when TAS1R1-TAS1R3 is expressed in other heterologous systems (26,27), supporting a role for TAS1R1-TAS1R3 signaling in STC-1 cells. Our data demonstrate a significant increase in umami taste receptor signal activity after pretreatment with butyrate. Of course, we cannot rule out that butyrate may modulate the expression of other genes involved in Ca^2+^signaling, such as Ca^2+^channels or other L-AA-sensitive GPCRs (8,39–41). However, it is still interesting to consider that butyrate exposure enhances L-Ala/IMP-induced Ca^2+^ signaling: this is a classical pathway associated with secretion of anorexergic gut hormones in EECs, which, in turn, elicit positive physiological effects, including blood glucose regulation and appetite reduction. Although butyrate alone does not induce gut hormone secretion, under conditions where a mixture of SCFAs are present, it may augment responses from other metabolites including propionate and thus will be interesting in future to see if there are alterations in taste receptor activity by propionate exposure (1,5,7,8,16,19,39).

SCFAs induce gut hormone secretion via signaling through their GPCRs.(12,16,18) Using the evidence gathered here, it is highly plausible that SCFAs also act to “reprogram” EECs to distinct, seemingly unrelated, dietary nutrients, by upregulating the counterparts receptive to their signaling. The temporal nature of these changes in terms of kinetics, and the duration in vivo will be important future steps to translate these findings. Despite this, our findings support the idea that GPCR signaling networks in EECs are highly complex, exhibiting the potential to adapt in response to dynamic fluctuations of bioactive nutrients (8,18,43,19–23,32,39,42). In summary, we can conclude that SCFA-induced remodeling of the GPCR signal system is an interesting and novel area that needs to be explored further, as it has potential therapeutic value.

## Author Contributions

M.S performed all experiments under supervision of E.W.T, G.F and A.C.H. M.S, A.J.B, G-A, E.W.T., G.F and A.C.H designed research and analyzed data and wrote the paper. All authors critically read and approved the final manuscript.

