## Supplemental Material for "Short chain fatty acids enhance expression and activity of the umami taste receptor in enteroendocrine cells via a Gα_i/o_ pathway"

### *Supplementary Material*

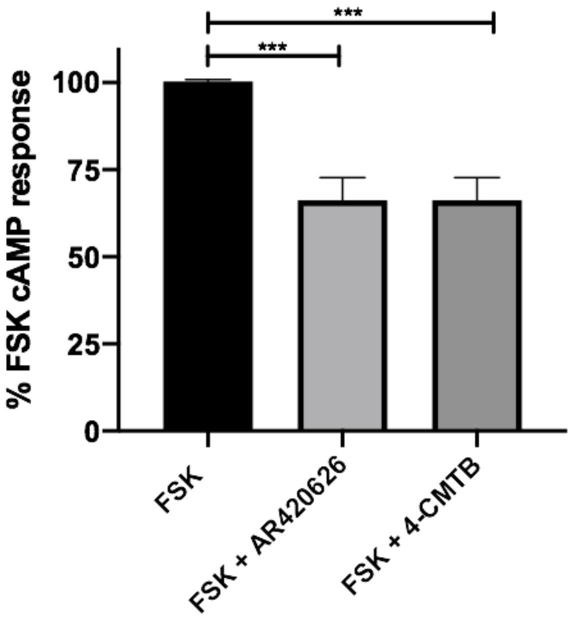


**Supplementary Figure 1.**

Intracellular cAMP accumulation measured in STC-1 cells on the addition of forksolin (3 µM) alone or with the addition of FFAR3-specific agonist AR420626 (10 µM) or FFAR2-specific agonist 4-CMTB (10 µM). Data is expressed as the percentage cAMP concentration compared to the FSK-only control (mean ± SEM; n=3); t-test *p<0.05; **p<0.01; ***p<0.001 vs. FSK-only control.

#### 1.1 Supplementary Tables

| Gene | Forward (5'-3') | Reverse (5'-3') |
|---|---|---|
| **α-gustducin** | TAGGAGCCGAGAGGACCAAG | GCTGGTATTCAGATGCCCTTTC |
| **β-actin** | GGCTGTATTCCCCTCCATCG | CCAGTTGGTAACAATGCCATGT |
| **FFAR2** | CGAGAACTTCACCCAAGAGC | TGAGGGAACTGAACACCACA |
| **FFAR3** | CAATACTCTGCATCTGTGAC | CAGGTAGACGGAAAAGAAA |
| **TAS1R1** | TGGAGGAGTGGTTGCGAAGA | TCCATGCCAACGTGAAGAGC |
| **TAS1R2** | TCCATGCCAACGTGAAGAGC | GCTACAGTTGTTGATTTCCTCCA |
| **TAS1R3** | TGCTGCCTACTGCAACTACAC | CCGGTCACTTAGCCGATCC |
| **TAS2R(108)** | TGACACGTCATTTGACCTCAG | GCTGGTCCTGTTTCTCTGCAT |
| **TAS2R(138)** | TCATTTCTGTTCCTTTCAGCCAT | CAGCAGTGCGATGTCACAGT |

**Supplementary Table 1.**

The forward and reverse primer sequences used for quantitative polymerase chain reaction (qPCR) analysis of the murine isoforms of house-keeping gene β-actin and the nutrient sensing receptors FFAR2, FFAR3, TAS1R1, TAS1R2, TAS1R3, TAS2R(108), TAS2R(138) and the gustatory G-protein α-gustducin
